# The BBS/CCT chaperonin complex ensures the localization of the adhesion G protein-coupled receptor ADGRV1 to primary cilia

**DOI:** 10.1101/2024.10.31.621306

**Authors:** Joshua Linnert, Deva Krupakar Kusuluri, Baran E. Güler, Sarita Rani Patnaik, Helen L. May-Simera, Uwe Wolfrum

**Author notes:** Corresponding author: Institute of Molecular Physiology, Molecular Cell Biology, Johannes Gutenberg University Mainz, Hanns-Dieter-Hüsch-Weg 17, 55128 Mainz, Germany. Authors contributed equally to this work. D. K. Kusuluri: Gena HealthX Private Limited, Business Development Centre, Rudolf-Diesel-Strasse 11, 69115 Heidelberg, GERMANY;, S. R. Patnaik: KAUST Smart-Health Initiative and Biological and Environmental Science and Engineering (BESE) Division, King Abdullah University of Science and Technology (KAUST), Jeddah, Saudi Arabia.

## Abstract

Primary cilia are antenna-like sensory organelles present on almost all eukaryotic cells. Their sensory capacity relies on receptors, in particular G-protein-coupled receptors (GPCRs) which localize to the ciliary membrane. Here we show that ADGRV1, a member of the GPCR subfamily of adhesion GPCRs, is part of a large protein network, interacting with numerous proteins of a comprehensive ciliary proteome. ADGRV1 is localized to the base of prototypic primary cilia in cultured cells and the modified primary cilia of retinal photoreceptors, where it interacts with TRiC/CCT chaperonins and the Bardet Biedl syndrome (BBS) chaperonin-like proteins. Knockdown of ADGRV1, CCT2 and 3, and BBS6 result in common ciliogenesis phenotypes, namely reduced ciliated cells combined with shorter primary cilia. In addition, the localization of ADGRV1 to primary cilia depends on the activity of a co-complex of TRiC/CCT chaperonins and the BBS chaperonin-like proteins. In the absence of components of the TRiC/CCT-BBS chaperonin co-complex, ADGRV1 is depleted from the base of the primary cilium and degraded via the proteasome. Defects in the TRiC/CCT-BBS chaperonin may lead to an overload of proteasomal degradation processes and imbalanced proteostasis. Dysfunction or absence of ADGRV1 from primary cilia may underly the pathophysiology of human Usher syndrome type 2 and epilepsy caused by mutations in *ADGRV1*.

## Introduction

Primary cilia are antenna-like organelles that can protrude from the cell membrane of most animal cells and receive external signals from the cellular environment(Mill et al., 2023). The microtubule pairs of their characteristic axoneme project from the mother centriole of the basal body, across the transition zone and into the cilia shaft of the cilia membrane. For their sensory function, primary cilia are equipped with numerous membrane receptors which convert the various external signals into intracellular signaling pathways (Satir and Christensen, 2007; Mill et al., 2023). Most of these receptors of primary cilia are G-protein-coupled receptors (GPCRs) localized to the ciliary membrane. Here, we focus on ADGRV1 in cilia, an adhesion G protein-coupled receptor (aGPCR) which is a subclass of G protein-coupled receptors (GPCRs). As seen in other aGPCRs, ADGRV1 features a unique combination of the GPCR characteristic seven-transmembrane helix domain (7TM) resembling the “receptor unit”, which transduces extracellular signals and mediates intracellular G-protein coupling via conformational changes, as well as an extracellular adhesion unit (Langenhan, 2020; Knapp et al., 2022). The extracellular N-terminal fragment (NTF) of aGPCRs can be released from the C-terminal fragment (CTF) by autoproteolytic cleavage at the GPCR proteolysis site in the GAIN (GPCR autoproteolysis-inducing) domain. This enables a tethered intramolecular agonist, the so-called “Stachel” sequence which comprises the first 5-10 amino acids of the CTF, to interact with the surface of the 7TM triggering the activation of aGPCRs (Scholz et al., 2019; Knapp et al., 2022; Liebscher et al., 2022). Interestingly, ADGRV1 can alternatively couple to two different heterotrimeric G-protein α−subunits (Shin et al., 2013; Hu et al., 2014) which may switch upon autoproteolytic cleavage and “Stachel”-mediated activation (Knapp et al., 2022).

Numerous mutations in the human *ADGRV1* gene are causative for the human Usher syndrome type 2 (USH2C) (Weston et al., 2004). USH is the most common form of combined hereditary deaf-blindness, characterized by both clinical and genetic heterogeneity. Up to four USH subtypes are related to defects in at least 11 genes (Fuster-García et al., 2021). Over the years, evidence has accumulated that USH could be classified as a ciliopathy, since USH molecules are often associated with primary cilia (Liu et al., 1997; Reiners et al., 2006; Maerker et al., 2008; Sorusch et al., 2019; Mill et al., 2023). In addition, more recently, mutations in *ADGRV1* have been associated with various forms of epilepsy (Wang et al., 2015; Myers et al., 2018; Liu et al., 2020; Leng et al., 2022; Zhou et al., 2022). Interestingly, there is recent emerging evidence, though, that defects in cilia are linked to the development of epilepsy (Karalis et al., 2022; Limerick et al., 2024; Vien et al., 2023).

The defects that most likely underlie USH2C are found in the sensory cells of the two sensory organs affected by USH, the auditory hair cells of the inner ear and the photoreceptor cells of the retina. Adhesion fibres, which are formed by the long extracellular domain of ADGRV1 and connect neighbouring membranes, are lost in Adgrv1-deficient mouse models (McGee et al., 2006; Michalski et al., 2007; Maerker et al., 2008). However, almost nothing is known about the pathomechanisms that lead to epilepsy associated with mutations in *ADGRV1*.

ADGRV1, previously named as very large GPCR 1 (VLGR1) is one of the largest receptor proteins of the human body with up to ∼6,300 amino acid residues and a molecular weight of ∼700 kDa (McMillan et al., 2002; McMillan and White, 2010; Hamann et al., 2015). ADGRV1 is almost ubiquitously expressed in mammals, with particularly high expression in the nervous system and the sensory cells of the inner ear and the eye (Güler et al., 2024; McMillan & White, 2010; Uhlén et al., 2015; https://www.proteinatlas.org/search/ADGRV1). We and others previously showed that ADGRV1 is part of various cellular adhesions: focal adhesions (Kusuluri et al., 2021; Güler et al., 2023), mitochondria-associated ER membranes (MAMs) (Krzysko et al., 2022), synapses between neurons, and neurons and glial cells (Reiners et al., 2005; Specht et al., 2009; Güler et al., 2024), ankle-links of developing hair cell stereocilia (McGee et al., 2006; Michalski et al., 2007) and the periciliary membrane complex at the base of the light-sensitive outer segment of vertebrate photoreceptor cells (Maerker et al., 2008).

Recent omics data and preliminary data on primary cilia in hTERT-immortalized retinal pigment epithelial cells (hTERT-RPE) indicated that ADGRV1 is a common molecular component of primary cilia (Knapp et al., 2022). We recently showed that ADGRV1 interacts not only with several other USH proteins but also with the CCT molecules of hetero-octameric TriC/CCT chaperonin complex and the three Bardet-Biedel syndrome (BBS) chaperonin-like proteins (BBS10, BBS12, and BBS6 also known as MKKS (McKusick-Kaufman-Syndrom) protein) in primary cilia (Linnert et al., 2023b) known to mediate the assembly of the BBSome at primary cilia (Seo et al., 2010; Zhang et al., 2012). However, whether ADGRV1 serves as a substrate of the TRiC/CCT/BBS chaperone co-complex remained unsolved.

Here, we identify potential interaction partners of ADGRV1 in the ciliary proteome and show a genetic/transcriptional interaction of *ADGRV1* and several genes of the primary ciliary core. We reveal that the absence of ADGRV1 and components of the TRiC/CCT/BBS chaperone co-complex leads to a common phenotype manifested in reduced ciliogenesis and ciliary length. Furthermore, we show that in the absence of the functional TRiC/CCT/BBS chaperone co-complex, ADGRV1 is depleted from the ciliary base by proteasomal degradation. Dysfunction or absence of ADGRV1 from primary cilia may underlay the pathophysiology of human Usher syndrome type 2 and specific forms of epilepsy caused by mutations in *ADGRV1*.

## Results

### Analysis of tandem affinity purification data revealed interactions of ADGRV1 with core proteins of primary cilia

To explore the general ciliary localization of ADGRV1, we co-immunostained ADGRV1 with the primary ciliary markers, anti-ARL13B or anti-GT335 and the nuclear DNA marker DAPI in primary human dermal fibroblasts, murine primary brain astrocytes and the mouse retinal photoreceptor cells (Figure 1). We could show by immunofluorescence microscopy that ADGRV1 is localized to the base of primary cilia of human fibroblasts, mouse astrocytes, and retinal photoreceptor cells (Figure 1).

**Figure 1.**
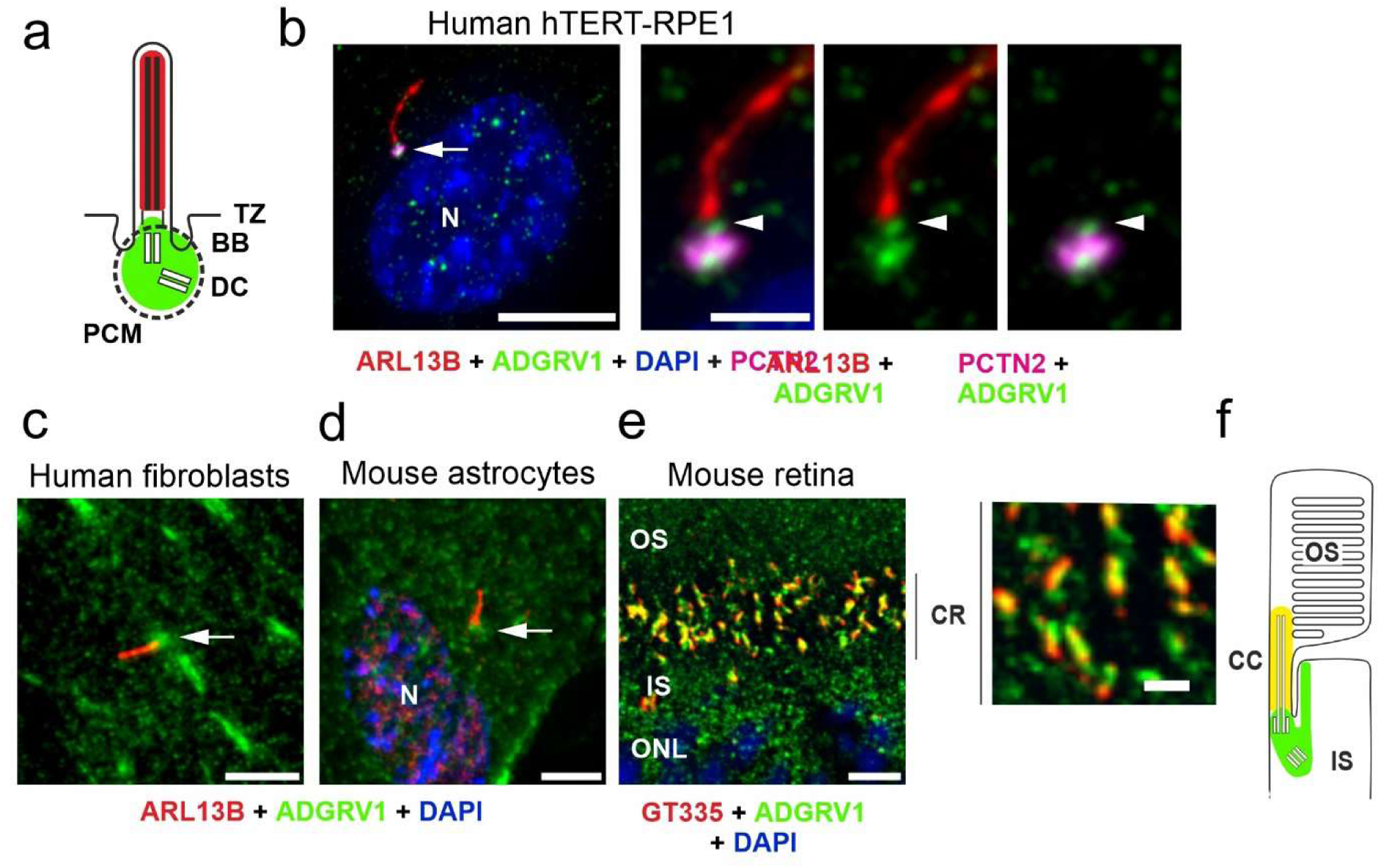
Conserved localization of ADGRV1 in primary cilia of various cell types. **(a)** Scheme of the prototypic primary cilium. The primary cilium is comprised of the ciliary membrane surrounding the microtubule-based axoneme, which is connected through the transition zone (TZ) to the basal body (BB, = mother centriole) and the adjacent daughter centriole (DC) surrounded by the pericentriolar material (dotted line). Green colour indicates localization of ADGRV1. **(b)** Trippel immunofluorescence of the axoneme marker ARL13B (red), the pericentriolar matrix marker pericentrin (PCTN2, magenta) and ADGRV1 (green) counter stained with the nuclear DNA marker DAPI in the primary cilium of hTERT-RPE1 cells. Staining pattern revealed the localization of ADGRV1 in the transition zone and the basal body complex (BB plus DC) surrounded by the pericentriolar material. **(c)** Localization of ADGRV1 in the primary cilium of human primary dermal fibroblasts and **(d)** in primary astrocytes derived from the hippocampus of a mouse brain. Double immunofluorescence of ARL1B (red) and ADGRV1 (green) counter stained with the DAPI revealed the localization of ADGRV1 at the ciliary base in both primary cell types. **(e)** Double immunofluorescence of the ciliary marker GT335 (red) and ADGRV1 (green) in longitudinal cryosection through the ciliary region (CR) of the photoreceptor cell layer of a mouse retina counter stained with DAPI. Staining pattern revealed the localization of ADGRV1 in CR where the connecting cilium (CC = transition zone of prototypic primary cilium) are localized at the joint between the inner segment (IS) and light sensitive outer segment (OS) of photoreceptor cells as indicated in (**f**) scheme of the ciliary region of a rod photoreceptor cell. Green colour indicates localization of ADGRV1; yellow: co-localization of ADGRV1 and GT335. Scale bars: a, left lower mag image = 10 µm and a, higher mag images, b, d, e = 5 µm

To determine potential molecular interactions of ciliary molecules with ADGRV1 we screened the existing datasets of mass spectrometry analyzed tandem affinity purifications (TAPs) with ADGRV1 constructs as baits (Figure 2a) (Knapp et al., 2022) for proteins of the ciliary proteome. For this, we compiled the cilia proteome by combining CiliaCarta (Van Dam et al., 2019) and the SYSCILIA gold standard v2 (Vasquez et al., 2021) identifying 1,178 ciliary proteins (Supplementary Table S1). The comparison of this ciliary proteome with the hits of the ADGRV1 TAPs revealed 116 proteins as putative interaction partners of ADGRV1 associated with primary cilia (Supplementary Table S2). Most of the potentially interacting proteins were observed in the TAP datasets of the N- and C-terminally tagged CTFs (81 and 69, respectively) and ADGRV1a (70) with a high degree of overlap (Figure 2b). This indicates that ciliary proteins preferentially interact with the 7TM domain of the receptor unit of ADGRV1. Subsequently, STRING analysis demonstrates that the identified potential interaction proteins are integrated into a large protein network (Figure 2c). These data have already been corroborated by interactions of ADGRV1 with a handful of these proteins (Reiners et al., 2005; van Wijk et al., 2006; Zou et al., 2017; Knapp et al., 2022; Linnert et al., 2023b).

**Figure 2.**
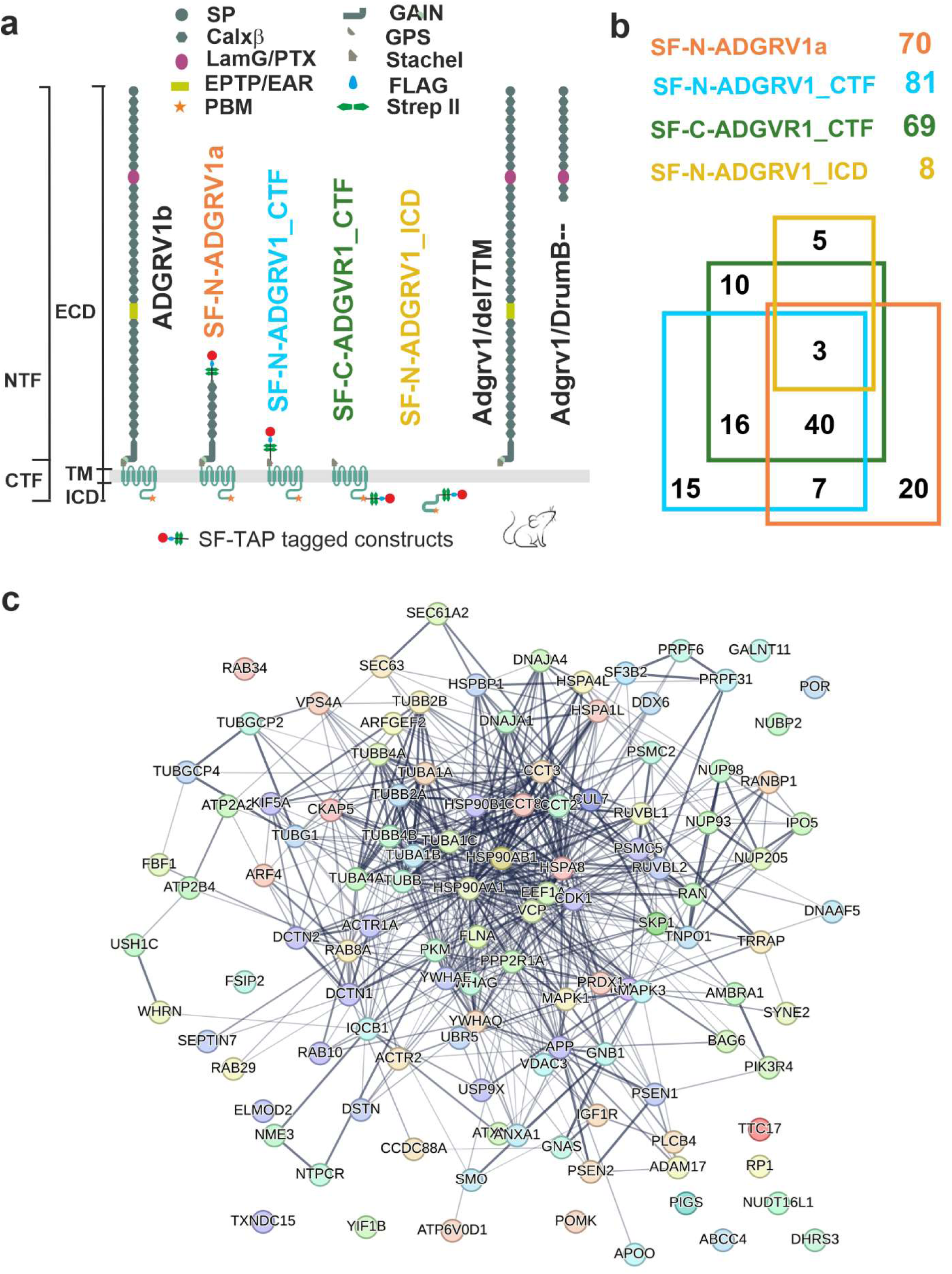
Interactions of ADGRV1 of proteins of primary cilium revealed by affinity proteomics. **(a)** Schematic representation of full length ADGRV1b, ADGRV1 bait constructs used in the tandem affinity purifications (Knapp et al., 2022), and truncated ADGRV1 proteins expressed in Adgrv1/del7TM and Adgrv1/DrumB-mouse models due to a deletion of the 7TM domain or a premature termination codon (McMillan & White, 2004; Potter et al., 2016). **(b)** Venn diagram of ADGRV1 prey assigned to the primary cilia proteome defined by the CiliaCarta and the *Syscilia* consortium (Van Dam et al., 2019; Vasquez et al., 2021). **(c)** The interaction of the prey, identified in b, visualised in a STRING network.

### Ciliary genes are differentially expressed in the transcriptomes of human USH2C patient-derived fibroblasts and the retina of *Adgrv1* deficient mice

We examined the effects of ADGRV1 deficiency on the expression of ciliary genes in cells and tissue previously obtained from dermal fibroblasts derived from a USH2C patient with the biallelic pathogenic variant *ADGRV1^Arg2959*^* compared to a healthy individual (Güler et al., 2024), and of retinae derived from the Adgrv1/del7TM mouse model compared to wild type (wt) control mice (Linnert et al., 2023a). We screened these transcriptomes for ciliary genes present in the combined datasets of CiliaCarta (Van Dam et al., 2019) and the SYSCILIA gold standard v2 (Vasquez et al., 2021) (Supplementary Table S1).

We identified 39 differentially expressed genes (DEGs) (17 upregulated and 22 downregulated), in USH2C patient-derived fibroblasts (Figure 3a). By applying STRING analysis, we found that the proteins corresponding to these genes organize in several distinct subclusters. (Figure 3b). Gene Ontology (GO) term analysis revealed that these ciliary DEGs were associated with the two *biological process* subcategories *G protein-coupled receptor signaling pathway* and *signaling* (Figure 3c; Supplementary Table S3).

**Figure 3.**
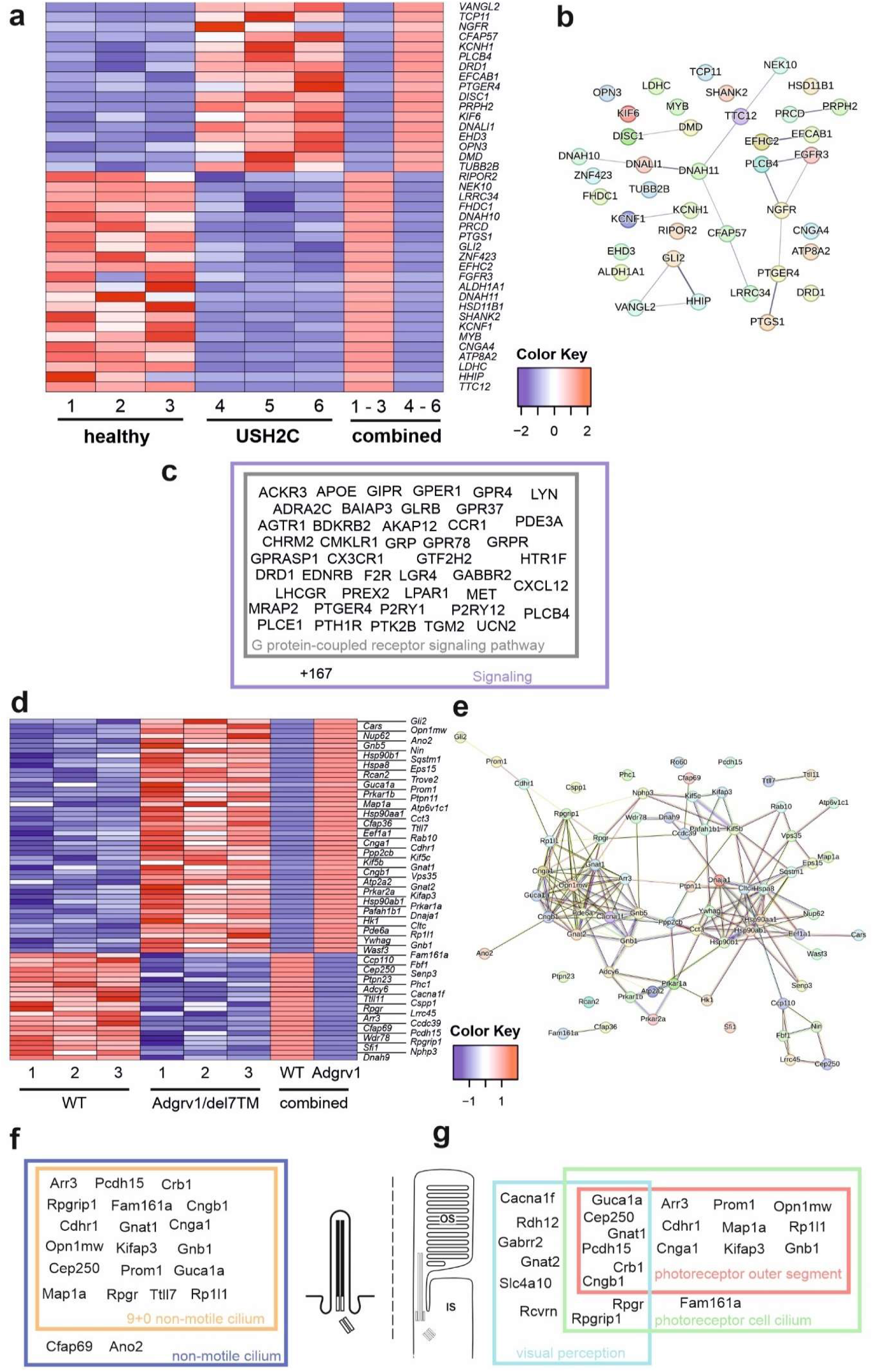
Differentially expressed ciliary genes in USH2C patient-derived fibroblasts and Adgrv1 deficient mouse retinae. (**a**) Heatmap of differentially expressed ciliary genes (DECGs) in patient-derived fibroblasts harbouring the p.R2959* nonsense mutation in the *USH2C* gene (USH2C) compared to fibroblasts derived from a healthy individual (healthy). (**b**) String network analysis of the interaction of the proteins corresponding to the DECGs identified in a. (**c**) Venn diagram of identified DECGs for the terms *G protein-coupled receptor signaling pathway* and *signaling* by a GO term analysis in category *biological process*. (**d**) Heatmap of DECGs in retinae of the Adgrv1/del7TM mouse model. (**e**) String network analysis of the interaction of the proteins corresponding to the DECGs identified in d. (**f,g**) Venn diagrams of identified DECGs for the terms *9+0 non-motile cilium and non-motile cilium* (f) and *photoreceptor outer segment, photoreceptor cell cilium* and *visual perception* (g) by a GO term analyses in category *cellular component*.

In the retina of Adgrv1/del7TM mice, 70 ciliary genes were differentially expressed (49 upregulated and 21 downregulated) (Figure 3d). STRING analysis revealed the proteins encoded by these ciliary DEGs into a protein-protein interaction network composed of two linked clusters (Figure 3e). Further analysis of these protein clusters showed their association with phototransduction and supporting proteins such as chaperones or vesicular transport proteins, respectively. GO term enrichment analysis revealed common links between the DEGs. In the category *cellular component,* we found 19 genes related to the term 9+0 non-motile cilium and 20 genes of the broader term *non-motile cilium*. Additionally, several genes were allocated to GO terms related to retinal photoreceptor cells, namely 14 genes to *photoreceptor outer segment*, 18 genes to *photoreceptor cell cilium* and 13 genes to *visual perception* (Figure 3f,g; Supplementary Table S4).

In summary, the absence of ADGRV1 resulted in dysregulation of the expression of genes associated with primary cilia and the highly specialized primary cilium of retinal photoreceptor cells.

### Ciliation and ciliary length is affected in human and murine primary cell models deficient for ADGRV1

We studied primary cilia in the following cellular models: human USH2C *ADGRV1^Arg2959*^*patient dermal fibroblasts (Figure 4) and mouse primary brain astrocytes derived from Adgrv1/del7TM and Adgrv1/DrumB mice (Figure 2a) in comparison to appropriate controls (Figure 5).

**Figure 4.**
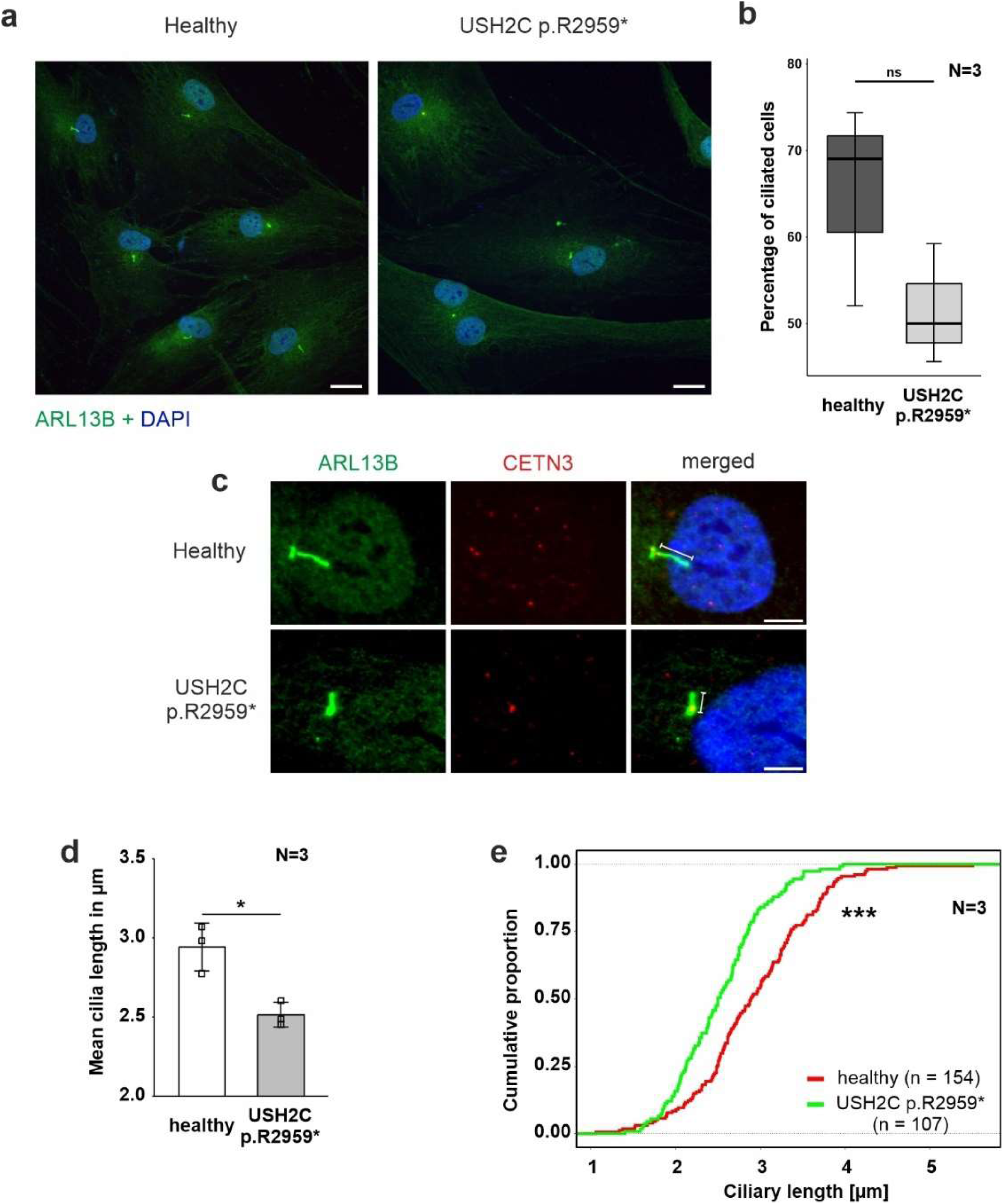
Phenotypic analysis of primary cilia of fibroblasts derived from a USH2C patient. (**a-e**) Immunofluorescence double staining of the axonemal marker ARL13B (green) and the ciliary base marker centrin 3 (CENT3, red) counterstained with nuclear DNA marker DAPI (blue) in patient-derived USH2C p.R2959* fibroblasts and healthy control cells. (**b,d,e**) Quantitative analysis of the number of ciliated cells, the mean cilia length and the distribution of cilia length in patient-derived fibroblasts and healthy control cells. Quantification revealed no significant changes in the number of ciliated cells, but significantly shorter cilia in the patient-derived fibroblasts compared to the healthy controls. N = number of biological replicates, n = number of analysed cells. Statistical significance was determined by the two-tailed Student’s t-test (b,d) and the Kolmogorov-Smirnov test (e): *p < 0.05, **p < 0.01, ***p < 0.005. Scale bar in a = 20 µm, c = 5 µm.

**Figure 5.**
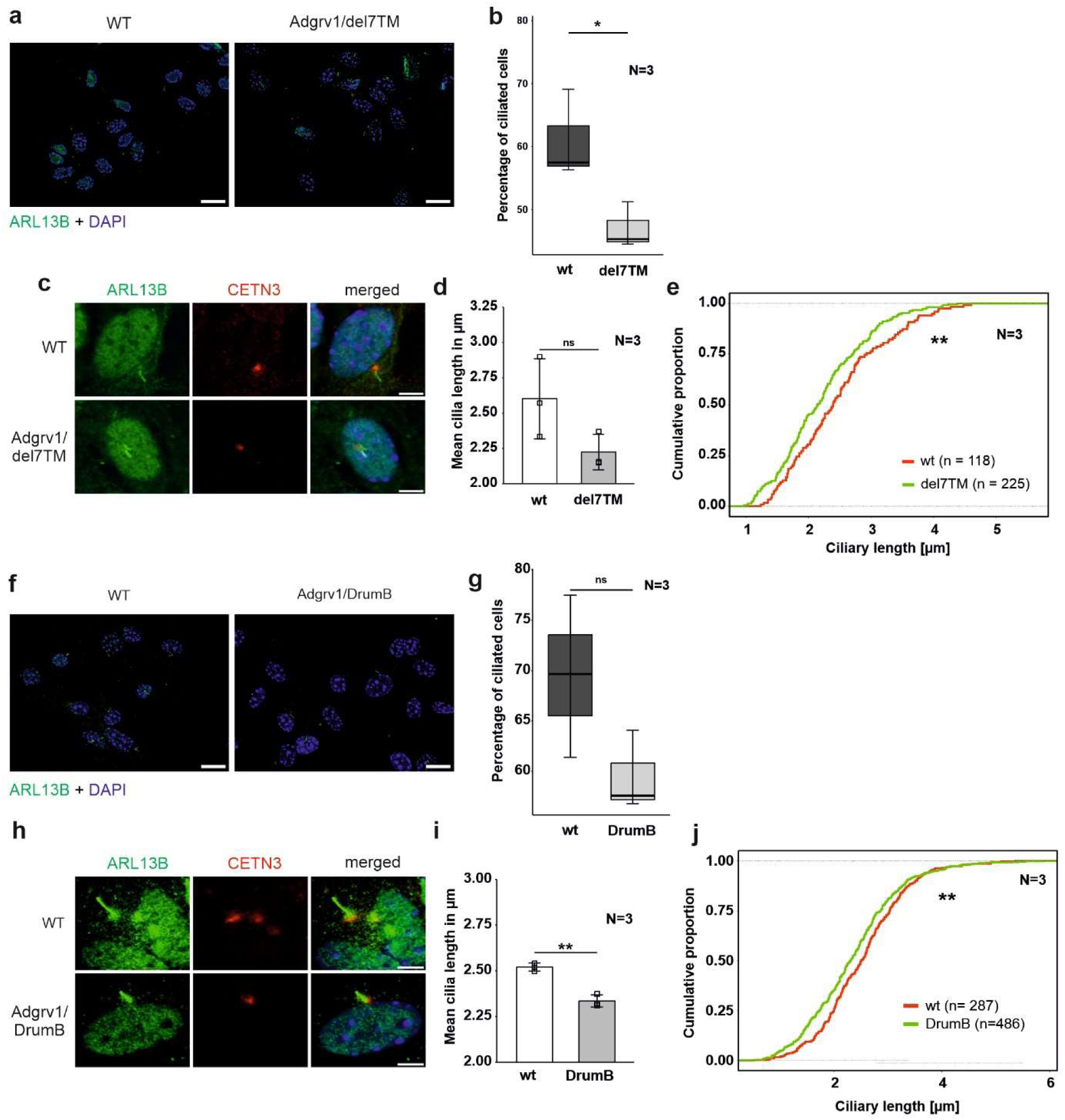
Phenotypic analysis of primary cilia of primary astrocytes isolated from brains of Adgrv1-deficent mouse models. (**a**) Immunofluorescence double staining of the axonemal marker ARL13B (green) and the ciliary base marker centrin 3 (CETN3, red) counterstained with nuclear DNA marker DAPI (blue) in primary astrocytes derived from Adgrv1/del7TM mice and wt controls. (**b-d**) Quantitative analysis of the number of ciliated cells (b), the mean cilia length (c) and the distribution of cilia length (d) of primary astrocytes revealed a significant decrease of ciliated cells in Adgrv1/del7TM astrocytes, no significantly altered mean cilia length, but a significant reduction of the cilia length quantifying the overall cilia length distribution. (**e**) Immunofluorescence double staining of the axonamal marker ARL13B (green) and the ciliary base marker centrin 3 (CETN3, red) counterstained with nuclear DNA marker DAPI (blue) in primary astrocytes derived from Adgrv1/DrumB mice and wt controls. Quantitative analysis of the number of ciliated cells (f), the mean cilia length (g) and the distribution of cilia length (h) of primary astrocytes revealed no significant decrease of ciliated cells in Adgrv1/DrumB astrocytes, but a significant reduction in the mean cilia length and the cilia length distribution. N = number of biological replicates, n = number of analysed cells. Statistical significance was determined by the two-tailed Student’s *t*-test (b,c,f-g) and the Kolmogorov-Smirnov test (d,h): *p < 0.05, **p < 0.01, ***p < 0.005. Scale bars: a, e, i = 5 µm.

Fibroblasts were starved for 48 hours in serum-free media to induce ciliation prior to immunohistochemical staining for the ciliary axoneme marker ARL13B and the ciliary base marker centrin 3 (Figure 4a). Fluorescence microscopy revealed reduced numbers of ciliated USH2C patient fibroblasts compared to healthy controls (Figure 4b). To determine the ciliary length, we measured the length of the ARL13B positive axoneme projecting from the centrin 3 positive ciliary base, calculated the mean of ciliary length (Figure 4c), and plotted the distribution of ciliary length (Figure 4c,d). Statistical evaluation of both analyses revealed a significant decrease in length in USH2C patient fibroblasts compared to healthy controls.

To analyze ciliogenesis in primary astrocytes we isolated astrocytes from the hippocampi of Adgrv1/del7TM, Adgrv1/DrumB, and wt control mice following our previously established protocol (Güler et al., 2021). Astrocytes were starved for 48 hours in serum-free media and stained for ciliary markers ARL13B and centrin 3 prior to fluorescence microscopic analysis (Figure 5). Fluorescence microscopy revealed reduced numbers of ciliated astrocytes derived from both the Adgrv1/del7TM and Adgrv1/DrumB mice compared to wt controls (Figure 5a, e). However, due to the high variability in the three independent experiments, statistical test revealed significantly reduced numbers of ciliated astrocytes only for Adgrv1/del7TM cells (Figure 5b,f).

Ciliary length measurements revealed shorter primary cilia of primary astrocytes isolated from both Adgrv1-deficient mouse models (Figure 5c,d,g,h). While in primary astrocytes of Adgrv1/DrumB mice both, the mean length and the length distribution of primary cilia were significantly different (Figure 5g,h) only the length distribution of primary cilia from Adgrv1/del7TM was significantly different (Figure 5c,d) compared to control.

Overall, we observed ciliary phenotypes in all ADGRV1-deficient cells tested, which manifested in the reduced number of ciliated cells and in a reduced length of primary cilia.

### Reduced length of the connecting cilium in photoreceptor cells of the Adgrv1/del7TM mouse retina

The light-sensitive outer segments of the photoreceptor cells of the vertebrate retina are highly modified primary sensory cilia characterized by an extended transition zone, named connecting cilium (Figures 1) (Roepman and Wolfrum, 2007; May-Simera et al., 2017). To determine whether the absence of ADGRV1 in photoreceptor cilia is also associated with a ciliary phenotype, we determined the length of the connecting cilium in the Adgrv1/del7TM mouse retina compared to wild-type mice. We immunostained the connecting cilium of photoreceptors in longitudinal retinal cryosections with antibodies against centrin 3 and GT335, both previously demonstrated as robust markers for the connecting cilium (Figure 6a,b). (Wolfrum, 1995; Grotz et al., 2022). Quantification of the mean cilia length and distribution revealed a significant reduction of connecting cilia length in the retinal photoreceptor cells of Adgrv1/del7TM mice when compared to those of wt mice (Figure 6c,d). This confirms the phenotype of reduced cilia length in Adgrv1-deficient photoreceptor cells observed in primary cell models, namely in dermal fibroblasts and brain astrocytes derived from USH1C patients and Adgrv1 mouse models.

**Figure 6.**
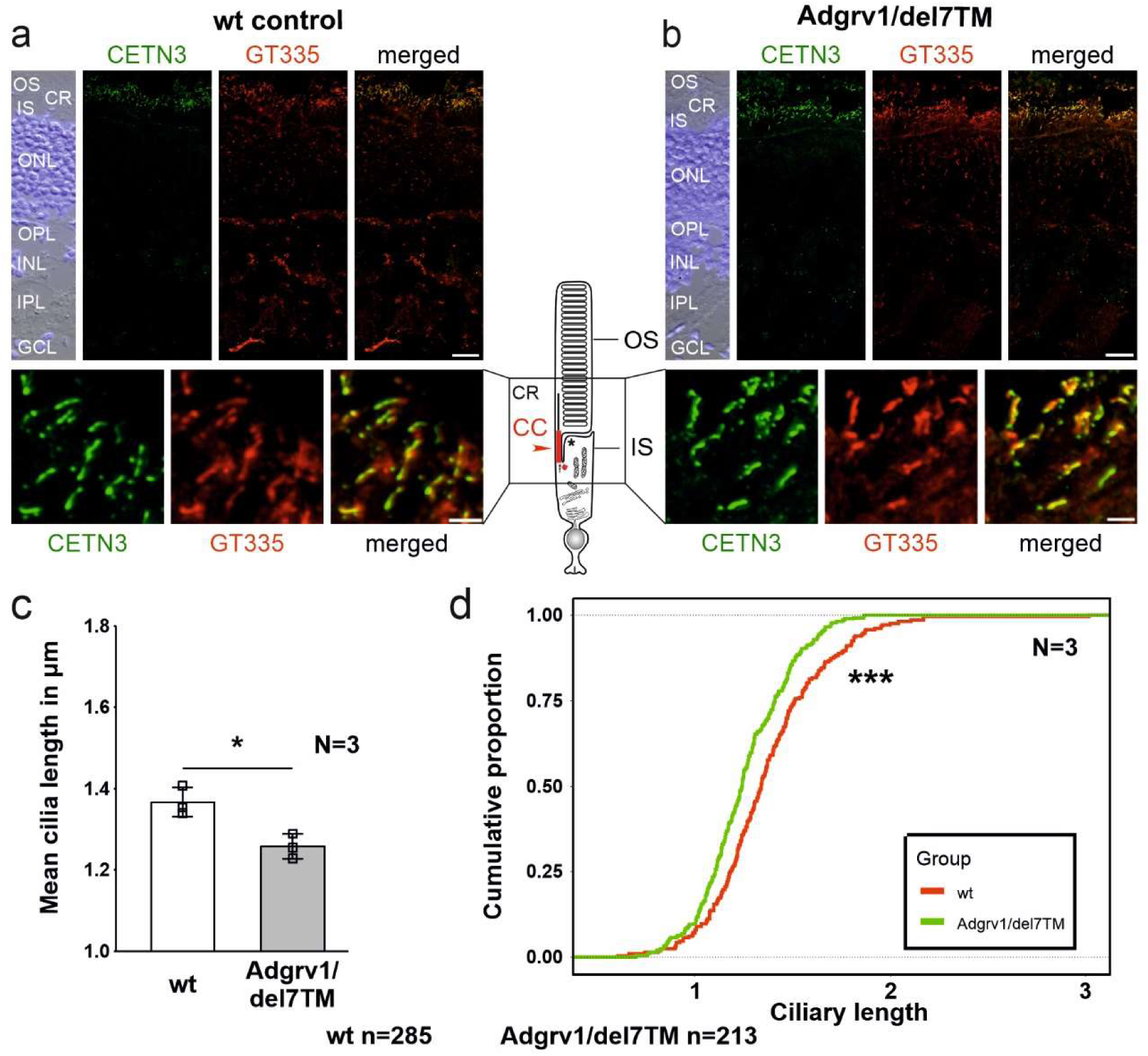
Primary cilium length in Adgrv1/del7TM retinae is reduced compared to wt controls. (**a,b**) Double immunofluorescence staining for centrin 3 (CETN3, green) and glutamylated tubulin (GT335) counterstained with DAPI in longitudinal cryosections through retinae of wt (a) and Adgrv1/del7TM mice (b). (**c**) Quantification of mean cilia length revealed a reduction in Adgrv1/del7TM retinae compared to wt controls confirmed by (**d**) the quantification of the distribution of the connecting cilia lengths. N = number of biological replicates, n = number of measured connecting cilia. Statistical significance was determined by the two-tailed Student’s *t*-test (c) and the Kolmogorov-Smirnov test (d): *p < 0.05, **p < 0.01, ***p < 0.005. Scale bars: a, b = 15 µm and 2 µm.

### The absence of TriC/CCT-BBS chaperonin complex components reduces ciliary localization of ADGRV1

We have previously shown that ADGRV1 physically interacts with components of the TRiC/CCT chaperonin complex and the chaperonin-like BBS molecules (Linnert et al., 2023b). Here we tested whether the localization of ADGRV1 at primary cilia depends on the TRiC/CCT-BBS chaperone complex, by depletion of chaperone complex components and immunostaining for ADGRV1 (Figure 7).

**Figure 7.**
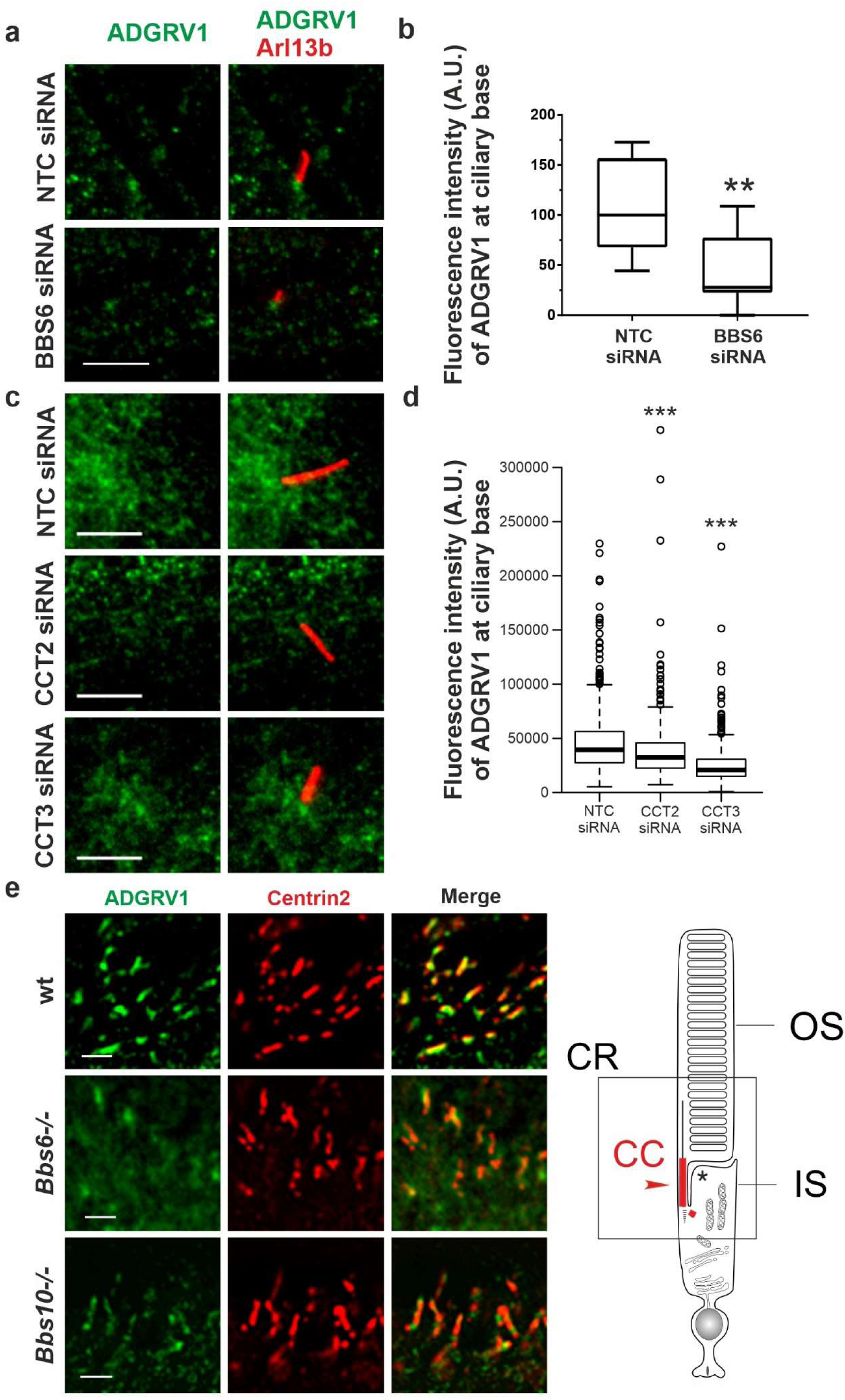
Immunofluoresence analysis of ADGRV1 in TriC/CCT-BBS chaperonin depleted cells and mouse retinae. **(a,c)** Double immunofluorescence staining for Arl13B (red) and ADGRV1 (green) of primary cilia in hTERT-RPE1 cells after siRNA-mediated knock down of (a) BBS6 and (b) CCT2 and CCT3 compared to non-targeting controls (NTC). (**b, d**) Quantification confirms reduced localization of ADGRV1 at the ciliary base upon BBS6 KD. (d) Quantification confirms reduced localization of ADGRV1 at the ciliary base upon CCT2 and CCT3 KD. **(e)** Double immunofluorescence staining for Adgrv1 (green) and connecting cilia marker Centrin2 (red) in longitudinal cryosections though the ciliary region (CR, in scheme left) of mouse photoreceptor cells revealed reduced Adgrv1 immunofluorescence in *Bbs6^-/-^*and *Bbs10^-/-^* mice when compared to wt littermate controls. Scheme shows CC: connecting cilium, OS: outer segment, IS: inner segment of a rod photoreceptor cell. Statistical significance was determined by the two-tailed Student’s *t*-test: *p < 0.05, **p < 0.01, ***p < 0.005. Scale bars: a,c = 5 µm; e = 1 µm.

For this, we depleted BBS6, CCT2, and CCT3 by siRNAs in hTERT-RPE1 cells (validation: Supplemental Figure S1), induced ciliogenesis by serum starvation and immunostained for ADGRV1 and the ciliary marker ARL13B (Figure 7). Fluorescence microscopy revealed that siRNA-mediated knockdowns of BBS6, CCT2, and CCT3 resulted in significant reductions of the intensity of the ADGRV1 immunofluorescence at the base of the primary cilium when compared to cells treated with non-targeting control (NTC) siRNAs (Figure 7a-d).

We also observed that the cilia length of the primary cilia was reduced after elimination of the three components of the TRiC/CCT/BBS chaperonin complex (Supplement Figures S2a-d). Interestingly, the knockdown of BBS6 did not alter the localization of CCT3 at the base of the cilium compared to that of the control cells (Supplemental Figure S3).

Next, we investigated whether the absence of BBS chaperonins also affects the distribution of ADGRV1 in the photoreceptor cilia (Figure 7e). For this, we immunostained Adgrv1 in the retinae of *Bbs6^-/-^* and *Bbs10^-/-^* knockout mice and a wt control Fluorescence microscopy of immunostained retinal longitudinal cryosections revealed prominent localization of Adgrv1 at the connecting cilium of photoreceptor cells of control wt retinae as revealed by counterstaining for the connecting cilium marker centrin 2 (Wolfrum, 1995; Giessl et al., 2004), coherent with previous findings (van Wijk et al., 2006; Liu et al., 2007; Maerker et al., 2008; Yang et al., 2010). In comparison, localization of Adgrv1 was reduced in the connecting cilia of photoreceptor cells in the *Bbs6^-/-^* and *Bbs10^-/-^*retinae (Figure 7e). Together, these findings indicate that the ciliary localization of ADGRV1 depends on the TriC/CCT-BBS chaperones.

### Depletion of BBS6 in hTERT-RPE1 cells leads to increased mRNA levels but decreased ADGRV1 protein expression

To validate whether the depletion of chaperonins affects the expression of *ADGRV1* we quantified the mRNA expression of *ADGRV1* upon BBS6 knockdown in hTERT-RPE1 cells (Figure 7). Quantitative real-time PCR (qPCR) analysis revealed a significantly elevated expression of *ADGRV1* after siRNA-mediated knockdown of BBS6, while the expression of *CCT3* was not altered (Figure 8a,b). Next, we analyzed the mRNA expression of *Adgrv1* in the retina of *Bbs6^-/-^* knockout mice. qPCR revealed that the expression of *Adgrv1* was significantly increased in postnatal 21 (P21) and P29 *Bbs6^-/-^*mouse retinae compared to wt littermates (Figure 8c,d), confirming the upregulation of the expression of *Adgrv1* mRNA transcripts in tissue.

**Figure 8.**
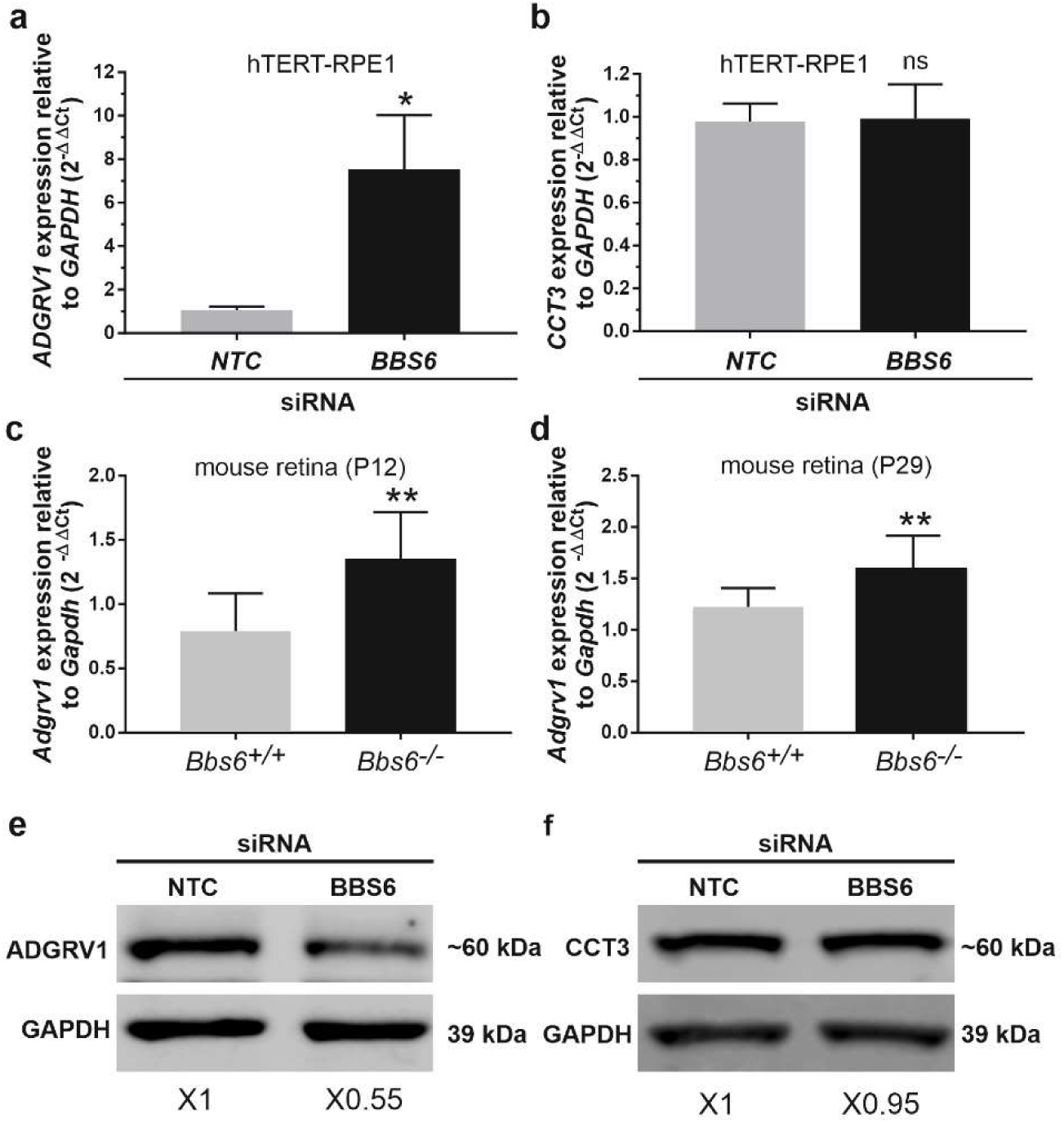
*ADGRV1* mRNA and ADGRV1 protein expression are changed in the opposite direction in cells and tissues deficient for BBS6. **(a)** Quantitative real-time PCR (qRT-PCR) analysis of *ADGRV1* mRNA expression in serum-starved hTERT-RPE1 cells after siRNA-mediated *BBS6* knockdown compared to non-targeting control (NTC). **(b)** qRT-PCR analysis of *CCT3* mRNA expression in serum-starved hTERT-RPE1 cells after siRNA-mediated *BBS6* knockdown compared to non-targeting control (NTC). Quantifications revealed an increase of *ADGRV1* but not of *CCT3* mRNA expression in *BBS6* depleted cells. **(c,d)** qRT-PCR analysis of *Adgrv1* mRNA expression in retinae of postnatal P12 and P29 *Bbs6^-/-^* mice demostrate significantly higher expression of *Adgrv1* mRNA. **(e,f)** Western blot analysis of ADGRV1 and CCT3 protein expression in serum-starved hTERT-RPE1 cells after siRNA-mediated *BBS6* knockdown. Quantifications of band densities revealed a decrease of ADGRV1 (x 0.55) but not of CCT3 expression (x 0.95) in *BBS6* depleted cells when compared to non-targeting controls (NTC). *GAPDH/Gapdh* and GAPDH protein were used as housekeeping control. Statistical significance was determined by the two-tailed Student’s *t*-test: *p < 0.05, **p < 0.01, ***p < 0.005.

To test whether the expression of the translated ADGRV1 protein is also affected by the depletion of BBS6 we performed Western blot analyses of lysates obtained from hTERT-RPE1 cells treated with siRNAs specific for BBS6 or NTC siRNA, respectively. Western blots probed with either antibodies against ADGRV1 or CCT3 revealed significantly reduced ADGRV1 protein expression, but no changes in the CCT3 protein expression in BBS6 knockdown cells (Figure 8e,f).

Taken together, in the absence of the BBS6 chaperonin ADGRV1 mRNA expression is upregulated while in contrast ADGRV1 protein levels are reduced. This suggests ADGRV1 expression is additionally post-translationally controlled or compensated at the protein level.

### Depletion of CCT/BBS chaperonins increases the proteasomal degradation of ADGRV1

To investigate whether the reduced ADGRV1 protein expression observed in the absence of chaperonins is due to proteasomal degradation, we experimentally inhibited the proteasomal pathway by MG132. For this, we treated BBS6 knockdown hTERT-RPE1 cells after ciliogenesis induction with MG132. Quantification of Western blots showed that MG132 treatments compensated the significant decrease of ADGRV1 protein expression following siRNA-mediated BBS6 knockdown (Figure 9a,b) indicating ADGRV1 is degraded via the proteasome in the absence of the BBS6 chaperonin.

**Figure 9.**
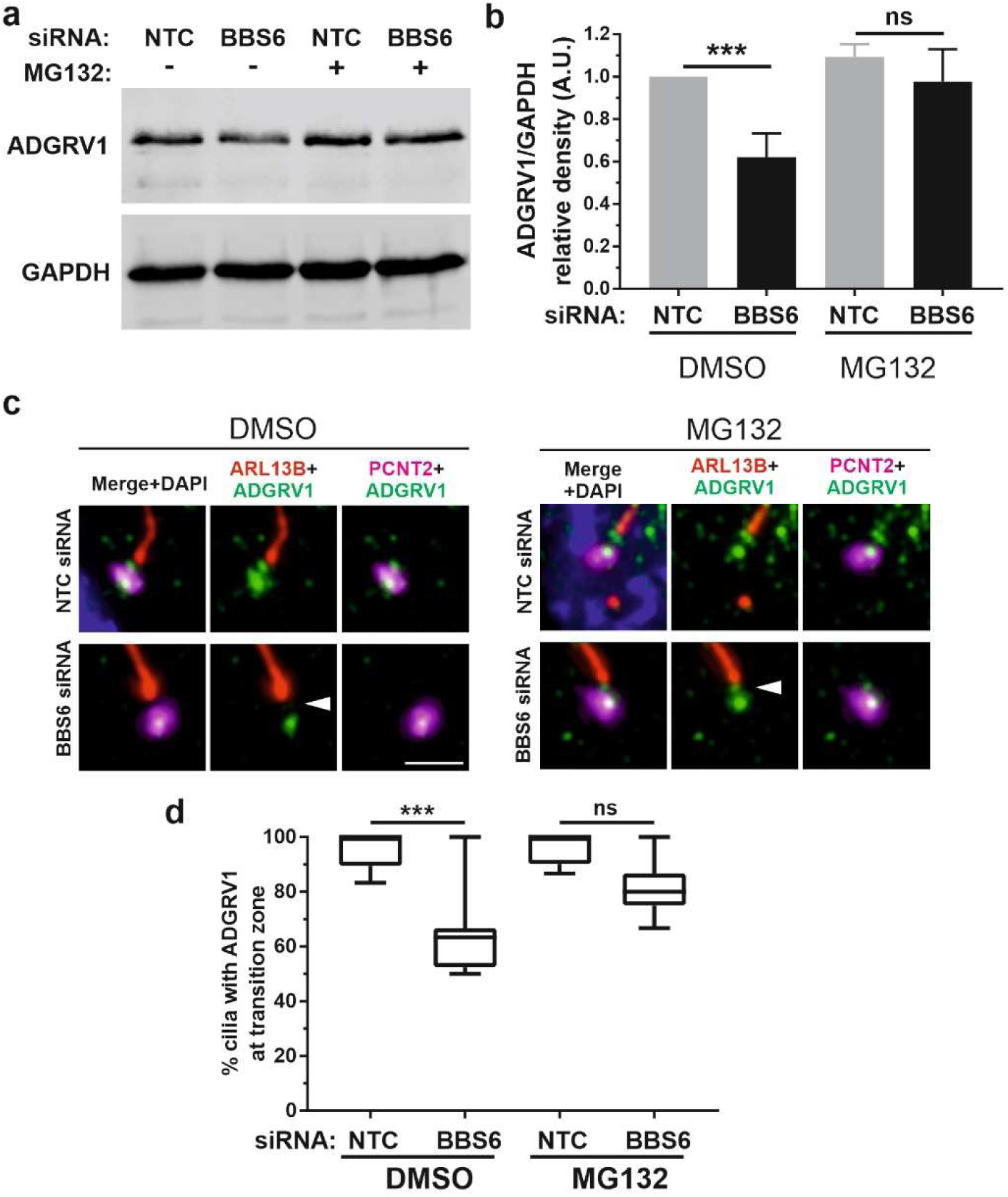
Increased proteasomal degradation of ADGRV1 upon BBS6 loss in hTERT-RPE1 cells. **(a,b)** Western blot analysis of ADGRV1 and CCT3 protein expression in serum-starved hTERT-RPE1 cells after siRNA-mediated *BBS6* knockdown treated with the proteasomal inhibitor MG132 solved in DMSO. The significant decrease of ADGRV1 protein expression in BBS depleted cells is restored by treatment with MG132. **(c,d)** Indirect immunofluorescence triple labelling of ADGRV1 (green) counterstained with axoneme marker ARL13B (red) and basal body marker pericentrin (PCNT2) (magenta) in NTC control and siRNA-mediated *BBS6* knockdown serum-starved hTERT-RPE1 cells treated with MG132 and DMSO. The ADGRV1 immunofluorescence in the transition zone of primary cilia (white arrowhead) is restored in cells treated with MG132. **(d)** Quantification of anti-ADGRV1 immunofluorescence intensity in the transition zone confirms recovery of ADGRV1 at the ciliary base and transition zone of BBS6 KD cells upon MG132 treatment. Statistical significance was determined by the two-tailed Student’s *t*-test: *p < 0.05, **p < 0.01, ***p < 0.005. Scale = 5 µm.

Next, we validated these findings *in situ* by immunocytochemistry (Figure 9c-f). For this we co-stained primary cilia of hTERT-RPE1 cells for the ciliary axoneme marker ARL13B, ciliary base marker pericentrin-2 (PCNT2), and ADGRV1. Fluorescent microscopy revealed the depletion of the anti-ADGRV1 fluorescence between the ciliary base stained by anti-PCNT2 and the axoneme stained by anti-ARL13B which represents the transition zone after BBS6 knockdown (Figure 9c). Quantification of ADGRV1 immunofluorescence confirmed that ADGRV1 protein remained in region of the ciliary transition zone when BBS6 knockdown cells were treated with the MG132. Together, this shows that ADGRV1 is prone to elevated proteasomal degradation in the absence of chaperonins.

## Discussion

Primary cilia are antenna-like cellular protrusions serving as receiver hubs coordinating various incoming environmental signals (Mill et al., 2023). These signals are sensed and transduced via GPCRs for subsequent transmission via downstream intracellular G protein-mediated signaling pathways (Hilgendorf et al., 2024). In the present study, we focus on the adhesion GPCR ADGRV1, its localization to the primary cilia and its role in ciliogenesis.

We identified several potential interacting proteins of the ADGRV1 interactome from TAP datasets within the comprehensive ciliary proteome, which we compiled from previously published datasets (Van Dam et al., 2019; Vasquez et al., 2021). In addition, our analysis of the differential transcriptomes of ADGRV1 deficient cells and tissues points towards diverse roles of ADGRV1 in cilia-related signaling pathways. Overall, the combined analyses of proteomic and transcriptomic data indicate that ADGRV1 functions as a receptor involved in primary cilia signaling.

During GPCR signal transduction cascades, ligand-induced conformational changes in activated GPCRs can couple to several different Gα subunits of heterotrimeric G proteins inducing different downstream signal pathways. Two alternative Gα subunits Gαi and Gαs have been previously described for ADGRV1 (Shin et al., 2013; Hu et al., 2014). More recently, we described that the activation of ADGRV1 by its autoproteolytic cleavage initiates the switch of G protein coupling from Gαi to Gαs (Knapp et al., 2022). Both α-subunits Gαi to Gαs have previously been described as GPCR signalling components in primary cilia (Hilgendorf et al., 2016). Accordingly, ADGRV1 can exert its dual function in primary cilia, firstly in adhesion, as found in the periciliary membrane complex of primary photoreceptor cilia (Maerker et al., 2008), and secondly in GPCR signal transduction and signaling coupled to G protein-mediated pathways. This hypothesis is supported by the differential transcriptomes obtained from the ADGRV1 deficient human USH2C patient fibroblast and the retina of the Adgrv1/delta7TM mouse model. GO term analysis of these differentially expressed genes demonstrated that the expression of 44 genes assigned to the term “*G protein coupled receptor signaling pathway*” is altered in the absence of ADGRV1.

However, there is evidence that ADGRV1 is also related to other non-GPCR signaling pathways such as Wnt-, sonic hedgehog (Shh)- and Notch-pathway (Knapp et al., 2022). Here, we found that the expression of genes coding for key components of these signaling pathways such as the Gli2 or Wnt ligands are dysregulated in the absence of ADGRV1. While these data suggest a potential involvement of ADGRV1 in Wnt, Shh, and Notch signaling, it is important to note that these findings are based primarily on dysregulated gene expression rather than direct mechanistic evidence. Future studies should focus on elucidating the precise molecular mechanisms linking ADGRV1 to these signaling pathways. Nevertheless, all these pathways have in common that they play important, vital roles in regulating primary cilia function and maintenance (Mill et al., 2023; Hilgendorf et al., 2024). The disruption of GPCR and non-GPCR mediated ciliary signaling pathways can lead to ciliary phenotypes such as impaired ciliogenesis and altered cilia length (Avasthi and Marshall, 2012; Canterini et al., 2017; Lee, 2020; Mukherjee et al., 2020). As observed in defective ciliary signalling pathways, mutations and deficiencies in ADGRV1 also lead to a reduced number of ciliated cells and shorter cilia which is in line with ADGRV1’s participation in signaling pathways of primary cilia.

ADGRV1 is localized to the base of primary cilia, namely to the transition zone and the periciliary lattice surrounding the basal body (present study, Knapp et al., 2022). In photoreceptor cilia, ADGRV1 is additionally found together with other USH proteins in the periciliary membranes (Maerker et al., 2008). There, it’s long extracellular domains form fibers that extend through the ciliary pocket and connect the cilia membrane of the transition zone to the periciliary plasma membrane. Overall, the ciliary base is a strategic compartment of the primary cilium that coordinates many essential ciliary functions. The transition zone contains the molecular machinery for sorting and controlling the molecular import and export, known as ciliary gating (Garcia-Gonzalo and Reiter, 2017; Park and Leroux, 2022; Wang et al., 2022). In addition, molecules at the ciliary base can regulate ciliary signaling, as recently shown for Shh signaling, which is positively regulated by Numb, a molecule associated with the endocytosis machinery at the ciliary pocket (Liu et al., 2024). Due to its localization ADGRV1 may participate in both, ciliary gating and the regulation of ciliary signaling. Interestingly, the latter function of ADGRV1 would be in line with the previously found association of ADGRV1 with endocytosis in the ciliary pocket (Bauß et al., 2014; Knapp et al., 2017) and the aforementioned link to the Shh signaling pathway.

Like other aGPCRs, ADGRV1 acts as a metabotropic mechanoreceptor sensing shear stress (Langenhan, 2020; Kusuluri et al., 2021). It is possible, that ADGRV1 is also activated by mechanical shear forces generated at the ciliary base and/or in the ciliary pocket by the bending of the ciliary shaft relative to the cell body. Consequently, the downstream signaling of activated ADGRV1 may regulate the functions described above.

At the base of primary cilia, ADGRV1 physically interacts with the components of the TRiC/CCT chaperonin complex and the three Bardet Biedl syndrome (BBS) chaperonin-like proteins BBS6, BBS10 and BBS12 (Knapp et al., 2022; Linnert et al., 2023b). Here, we demonstrate that ADGRV1 also functionally interacts with these chaperonins in primary cilia. siRNA-mediated knockdowns of ADGRV1 and CCT2 molecules in hTERT-RPE1 cells lead to a consistent ciliary phenotype, namely altered ciliogenesis paired with reduced ciliary length, as previously reported for BBS chaperonin-like proteins (Hernandez-Hernandez et al., 2013; Patnaik et al., 2019; Hey et al., 2021) indicating that ADGRV1 and the TRiC/CCT-BBS chaperonins are enrolled in overlapping ciliary pathways. Furthermore, we show that the ciliary localization of ADGRV1 at the ciliary base depends on the presence of the chaperonin proteins BBS6, CCT2 and CCT3. ADGRV1 protein is depleted from the ciliary base of primary cilia in cultured cells and from the photoreceptor cilium in retinal tissue sections upon loss of these components. These findings were confirmed by determining the amount of ADGRV1 in protein lysates of BBS6 knockdown cells. However, the absence of BBS6 does not lead to a reduction in transcriptional activity of the *ADGRV1* gene; in fact, *ADGRV1/Adgrv1* mRNA levels were several-fold higher after knockdown *BBS6* or knockout of *Bbs6* in cells and retinal tissue, respectively. The upregulation of *ADGRV1* transcription could be part of a compensatory mechanism that responds to the reduced amount of ciliary ADGRV1 protein. There is evidence that BBS6 regulates the nuclear import of SMARCC1, a chromatin remodeler that modifies gene activity in the developing heart (Scott et al., 2017). However, whether a function of BBS6 in cytoplasmic-nuclear shuttling also plays a role in the transcriptional regulation of ADGRV1 gene activity needs to be clarified in future studies. In any case, treatments with the proteasomal inhibitor MG123 restore protein expression, demonstrating that the ADGRV1 protein is degraded by the proteasome in the absence of chaperonins.

Substrates for the CCT chaperonin complex are predominantly cytoplasmic proteins such as the cytoskeletal components actin and tubulin as well as several proteins with β-propellers/WD40 repeats (Yam et al., 2008; Vallin and Grantham, 2019). At the base of primary cilia, the CCT chaperonins and BBS chaperonin-like proteins form a CCT/BBS chaperonin complex which associates with a subset of BBS molecules to mediate the assembly of the BBSome (Seo et al., 2010; Zhang et al., 2012; Wingfield et al., 2018). Screens in yeast also demonstrated membrane proteins as putative substrates of CCT chaperonins (Dekker et al., 2008) which is in line with our finding that transmembrane receptor ADGRV1 is a substrate of the CCT/BBS chaperonin complex. We hypothesize that the function of the CCT/BBS chaperonin complex ensures correct ADGRV1 conformation for proper integration of the receptor at the ciliary base. If this is not the case, misfolded ADGRV1 molecules are transferred to the proteasome for proteolytical degradation. This proteostasis pathway is well known from the, where misfolded GPCRs are targeted to the proteasome (Meusser et al., 2005). As found for defective of the quality control system in the ER, defects in CCT/BBS chaperonin system may lead to an overload of proteasomal degradation processes for a prolonged period and to imbalanced proteostasis (Pavel et al., 2016; Álvarez-Satta et al., 2017; Faber and Roepman, 2019).

Mutations in the genes for *ADGRV1*, *MKKS/BBS6* and *CCT2* are causative for severe syndromic retinopathies, namely USH2C, BBS, and MKKS (Sheffield, 2010; Slavotinek, 2020; Fuster-García et al., 2021) or non-syndromic retinal dystrophies such as Leber congenital amaurosis (LCA) (Minegishi et al., 2016; Weihbrecht et al., 2017). The proteins encoded by these genes are colocalized and likely involved in common processes. It is therefore plausible that the pathomechanisms leading to retinopathies overlap and have a common origin at the photoreceptor cilium. Our findings also raise the possibility of a common treatment target for these diseases, namely photoreceptor proteostasis. In animal models for autosomal dominant Retinitis pigmentosa due to mutations in the *RHO* gene, recent preclinical findings indicate that treating imbalanced proteostasis by pharmacological inhibition of the VCP/proteasome rescues photoreceptor degeneration (Sen et al., 2021). These findings suggest treatment strategies for some forms of USH, BBS and LCA.

Our findings align with the broader hypothesis that defects in primary cilia structure or signaling contribute to neurological conditions such as epilepsy (Karalis et al., 2022). Mutations in human *CILK1* (ciliogenesis-associated kinase 1) have been linked to both ciliopathies and epilepsy (Limerick et al., 2024) which is consistent with the disease manifestation resulting from mutations in *ADGRV1*. As in the case of pathogenic *ADGRV1* mutations the ciliation rate, the ciliary length, and Shh signaling are altered by *CILK1* mutations. Although there is growing evidence that dysfunctions of primary cilia are associated with epilepsy (Karalis et al., 2022; Vien et al., 2023; Limerick et al., 2024), it is not clear to what extent these are caused by defects in ADGRV1. Whether the dysregulation of glutamate homeostasis in hippocampal astrocytes of the brain of ADGRV1-deficient mice epilepsy (Güler et al., 2024) is associated with cilia defects needs to be determined in future studies.

However, it remains unclear whether the epileptic phenotypes observed in ADGRV1-deficient models are primarily due to ciliary dysfunction or if secondary effects, such as altered glutamate homeostasis, play a more prominent role.

## Conclusion

Present data indicate that the aGPCR ADGRV1 is part of the ciliary protein interactome and participants in ciliary signalling pathways. At the ciliary base ADGRV1 interacts with the TRiC/CCT-BBS chaperonin co-complex which ensures its correct ciliary localization. Failure of the chaperonin complex leads to proteasomal degradation of ADGRV1 which may result in imbalanced proteostasis in photoreceptor cells. Dysfunction or absence of ADGRV1 from primary cilia may underlay the pathophysiology of human USH type 2 and epilepsy caused by mutations in *ADGRV1*.

## Material and Methods

### Animals and tissues

Animals were used in accordance with the guidelines set by the Association for Research in Vision and Ophthalmology. The use of mice for research purposes was approved by the local authority, Mainz-Bingen district administration under file number 41a/177-5865-§11 ZVTE on April 30, 2014.

Laboratory mice were housed under 12/12-h light/dark cycles, food and water ad libitum. Following mouse models were: Adgrv1/del7TM mice express only a truncated form of the Adgrv1 protein lacking the entire 7-transmembrane and intracellular domain, due to the replacement of 101 bp in exon 82 with a vector containing premature termination codon (McMillan & White, 2004). Adgrv1/DrumB mice carry a nonsense mutation (c.8554+2t>c) in intron 37-38 of Adgrv1, leading to a premature termination codon (Potter et al., 2016). *Mkks/Bbs6^-/-^* knockout mice have been previously described by Ross et al. (2005). Breeding background of the mice was the C57BL/6 strain.

Brain tissue, eyeballs and isolated retinae were removed from the sacrificed animals for subsequent experimental protocols. Eyeballs and retinal tissue from *Bbs10^-/-^* and *Bbs12^-/-^* complete knockout mice (Cognard et al., 2015) were received from Dr. Vincent Marion (Institut national de la santé et de la recherche médicale, Strasbourg, France).

### Antibodies

Following primary antibodies were used: mouse anti-GT335 (AdipoGen, AG-20B-0020), rabbit anti-ARL13B (Proteintech, 17711-1-AP), mouse anti-ARL13B (Proteintech, 66739-1-Ig), mouse anti-CETN3 (Giessl et al., 2004), rabbit anti-CETN3 (Giessl et al., 2004), rabbit anti-ADGRV1 (Reiners et al., 2005), goat anti-CETN2 (Giessl et al., 2004), rabbit anti-BBS6 (Proteintech, 13078-1-AP), mouse anti-GAPDH (Abcam, ab9484), mouse anti-CCT3 (Proteintech, 60264-1-Ig), goat anti-PCTN2 (Santa Cruz Biotechnology, sc-28145). Secondary antibodies were conjugated to Alexa 488, Alexa 555, Alexa 568, Alexa 680, or IR Dye 800, all purchased from Invitrogen or Rockland Immunochemicals. Nuclear DNA was counterstained with DAPI (40,6-diamidino-2-phenylindole, 1 mg/ml, diluted 1:12 000; Sigma-Aldrich).

### Cell Culture

hTERT-RPE-1 cells: hTERT-immortalized retinal pigment epithelial cells (RPE) were cultured in Dulbecco’s Modified Eagle Medium/Nutrient Mixture F-12 (DMEM/F-12) (Thermo Fisher Scientific).

HEK293T cells: Human HEK293T cells were cultivated in Dulbecco’s Modified Eagle Medium (DMEM) (Thermo Fisher Scientific) containing 10% heat-inactivated foetal calf serum (FCS). For transient transfections, GeneJuice^®^ (Merck Millipore) or Lipofectamine LTX with Plus reagent (Thermo Fisher Scientific) were used according to the manufacturer’s instructions.

Human primary fibroblasts: Healthy primary fibroblast lines were expanded from skin biopsies collected from human donors (ethics vote: Landesärztekammer Rhineland-Palatinate to Kerstin-Nagel Wolfrum). ADGRV1 R2959* patient-derived fibroblasts were a kind gift from Dr. Erwin van Wijk (Radboud University Medical Center, Nijmegen) and were derived from skin biopsies of a 57-year-old male USH2C patient who carries a nonsense mutation in the VLGR1/ADGRV1 gene (g.[90006848C>T]) (Usher syndrome database, https://databases.lovd.nl/shared/variants/GPR98/unique). The primary human dermal fibroblast lines were mycoplasma negative and cultured in DMEM, containing 10% FCS and 1% penicillin-streptomycin at 37°C and 5% CO_2_.

To induce growth of primary cilia, cells were cultured for 48 h in serum-free OptiMEM (Thermos Fisher Scientific).

### Proteasome inhibition

For the analysis of the proteasome activity in BBS6/CCT2/CCT3 depleted hTERT-RPE1 cells, the proteasome was inhibited by a treatment with MG132 (Sigma). For ciliation experiments, cells were seeded on glass coverslips (15,000 cells). The following day the complete growth media was substituted with serum-free OptiMEM (Thermos Fisher Scientific). To induce growth of primary cilia, cells were starved for 48 h. 5 µM MG132 was added to the cells in the last two hours. Control cells were treated with an appropriate volume of DMSO.

### siRNA-mediated knockdown

hTERT-RPE1 cells were transfected with siRNAs using Lipofectamine RNAiMAX (Invitrogen, Thermo Fisher Scientific) according to the manufacturer’s instructions. siRNAs were used for following human genes: *BBS6* (L-013300-00-0005, Dharmacon)*, CCT3* (hs.Ri.CCT3.13.1, IDT)*, CCT2* (hs.Ri.CCT2.13.1, IDT).

### Isolation and culturing of primary murine astrocytes

Hippocampal astrocytes were isolated from newborn mouse pups on postnatal day 0 (PN0) as previously described (Güler et al., 2021). Astrocyte cultures of Adgrv/del7TM, Adgrv1/DrumB^-/-^ and C57BL/6 were cultivated in DMEM, containing 10% FCS and 1% penicillin-streptomycin at 37°C and 5% CO_2_.

### Immunocytochemistry of primary cilia

For immunostaining, cells were cultured in 24-well plates on coverslips. Cells were fixed in PBS containing 4% paraformaldehyde for 10 min at room temperature (RT). After fixation cells were washed three times with PBS and permeabilised by using PBST (0.2% Triton-X-100, Roth) for 10 min at RT, washed once with PBS, and incubated for 1 h in blocking solution blocked (0.1% ovalbumin, 0.5% fish gelatine in PBS) at RT. Primary antibodies in blocking solution were incubated overnight at 4°C. Subsequently, cells were washed three times with PBS prior to the incubation with secondary antibodies, containing DAPI, for 1 h at RT. After three washes with PBS, cells were mounted with Mowiol (Roth) prior to microscopic analyses.

### Immunohistochemistry

Dissected eye cups were cryofixation in melting isopentane (-168°C) as previously described (Wolfrum, 1991). Cryofixed specimens were cryosectioned in a MICROM HM 560 cryostat (Thermo Scientific). 10-12 µm thick cryosections were placed on 0.01% poly-L-lysine-coated coverslips and encircled by hydrophobic barrier pap pen (Science Services). Subsequently, cryosections were incubated subsequently with 0.01% Tween 20 for permeabilisation, washed several times, and incubated for 45 min in blocking solution at RT. Primary antibodies were incubated overnight at 4°C and after washing, secondary antibodies including DAPI (1 mg/ml, Sigma) were incubated for 1h at RT. After several washes, sections were mounded in Mowiol (Roth) and stored at 4°C in the dark.

### Western blot analysis

Western blotting was performed as previously described (Krzysko et al., 2022). In brief, protein lysates from cells and tissue were prepared using Triton-X-100 lysis buffer (50 mM Tris/HCl, 150 mM NaCl, 0.5% Triton-X-100, pH 7.4) containing complete protease inhibitor cocktail (Sigma) prior to sonication. Protein content was determined by performing a BCA protein assay (Merck Millipore). SDS-PAGE was performed on a Mini-PROTEAN System (BioRad) and separated proteins were transferred to a polyvinylidene difluoride (PVDF) membrane (Merck Millipore) using the Mini Trans-Blot Electrophoretic Transfer Cell (BioRad). Blot membranes were blocked in AppliChem blocking reagent for 1 h at RT. Primary antibodies were diluted in blocking solution and incubated overnight at 4°C prior to the incubation of secondary antibodies, diluted in blocking solution, for 1 h at RT. Blot membranes were scanned with the Odyssey infrared imaging system (LI-COR Biosciences).

### Microscopy, image analysis, and data processing

Mounted cells and tissue were analysed on a Leica DM6000B microscope and digital images were obtained by using a Leica K5 digital camera (Leica) and stored as LIF-files (Software Leica Application Suite). Microscopic images were processed using the Fiji/ImageJ software(Schindelin et al., 2012). For the measurement of primary cilia length, images were loaded into Fiji using the Bio-Formats plugin. Primary cilia were identified by the markers for the ciliary axoneme and the basal body.

Ciliary length was measured by applying the Fiji line draw tool. Quantification of anti-ADGRV1 signal at the primary cilia base was measured and calculated by the total corrected cellular fluorescence (TCCF = integrated density- (area of selected cell x mean fluorescence of background readings) (McCloy et al., 2014). The cilia base area was determined by a counterstaining with the ciliary base marker anti-PCTN2. The PCTN2 positive area was then used to measure the anti-ADGRV1 signal.

For densitometry analysis of Western blots, the LI-COR software Image Studio was used. For quantification, we normalised the detected bands to the bands of the housekeeping protein GAPDH.

Statistical analysis and figure preparation was conducted with RStudio (R Core Team, 2022). To determine the degree of colocalization of the immunostaining for ADGRV1 and Centrin 2 in mouse retinae the Pearson correlation coefficient was used (Adler and Parmryd, 2010). Pearson coefficient was calculated using the Coloc 2 plugin of ImageJ/Fiji (https://imagej.nih.gov/ij/). The correlation value has a range from +1 to -1. A value of 0 indicates no association, while a value greater than 0 indicates positive association and a value less than 0 indicates negative association.

### Reverse transcriptase (RT)-qPCR

Total RNA was isolated from hTERT-RPE1 cells and mouse retinae according to the instructions of the Qiagen RNeasy Mini Kit. RNA quality was determined by a Nanodrop (Thermo Fisher Scientific). RNA was transcribed to cDNA according to the instructions of the SuperScript™ III First-Strand Synthesis SuperMix (Thermo Fisher Scientific). Subsequently, qPCR was performed using a QuantStudio 1 qPCR machine (Thermo Fisher Scientific) and the iTaq Universal SYBR Green Supermix (Bio Rad). Following primers were used: Human: *CCT2* fwd 5’ TAAGGCAGGAGCTGATGAAG 3’; rev 5’ TTTAGCTGCTGGATTGTCAAC 3’; *CCT3* fwd 5’ ATTGTGCTGCTGGATTCTTCTCTG 3’; rev 5’ CTGCGGATGGCTGTGATATTGG 3’; *BBS6* fwd 5’ AATGACACTGCCTGGGATG 3’; rev 5’ TCGTTGTGAGTCTTGTGTCTG 3’; *GAPDH* fwd 5’ GAGTCAAGGGATTTGGTCGT 3’; rev 5’ TTGATTTTGGAGGGATCTCG 3’. Mouse: *Gapdh* fwd 5’ CGACTTCAACAGCAACTCCCACTCTTCC 3’; rev 5’ TGGGTGGTCCAGGGTTTCTTACTCCTT 3’; *Bbs6* fwd 5’ GTCTGCTCTGCAAGATTTGG 3’; rev 5’ AAGACGTGCATTGCTGTTTGG 3’; *Adgrv1* fwd 5’ CCGATGGTGCCAGATATAAAGTAG 3’; rev 5’ TCTTCGCTTCCTGGAGAATTG 3’

### Analysis of TAP and RNAseq data

With the datasets, differential gene expression analysis and GO enrichment analysis were performed (https://en.novogene.com/services/research-services/transcriptome-sequencing/mrna-sequencing/). Databases for GO term analysis and STRING networks of RNAseq data were accessed on 12.06.2024. STRING database for the TAP data was accessed on 13.03.2024.

## Supporting information

Supplementary Table S1 Compiled ciliary proteome

Supplementary Table S2 Ciliary proteins in TAP data sets

Supplementary Table S3 GO term USH2C fibroblasts

Supplementary Table S4 GO terms Adgrv1 retina

## Acknowledgements

We thank Ulrike Maas, Yvonne Kerner, and Gabi Stern-Schneider for excellent technical assistance, Dr. Kerstin Nagel-Wolfrum for critical discussion of the present data, Dr. Vincent Marion for kindly providing tissue from Bbs10-/- and Bbs12-/- mice, and Dr. Erwin van Wijk for kindly providing ADGRV1 R2959* patient cells.

## Supplementary Materials

Supplementary Table S1: Compiled ciliary proteome

Supplementary Table S2: Ciliary proteins in TAP data sets

Supplementary Table S3: Gene Ontology (GO) term analysis of DEGs in USH2C patient-derived fibroblasts

Supplementary Table S4: Gene Ontology (GO) term analysis of DEGs in Adgrv1/delTM mouse retinae

## Data availability statement

The mass spectrometry proteomics data have been deposited to the ProteomeXchange Consortium via the PRIDE partner repository with the dataset identifier PXD042629. The RNA-seq data presented in this study has been deposited in the NCBI Sequence Read Archive (SRA). It is accessible under the BioProject accession numbers: PRJNA1180279 (Adgrv1/del7TM retinae, WT retinae) and PRJNA1180475 (USH2C patient-derived fibroblasts, healthy donor fibroblasts).

## Author contributions

All authors contributed to the article and approved the submitted version. D.K.K. and J.L. contributed to the TAP data analyses, immunocytochemistry, ciliogenesis experiments, and protein-protein interaction assays. J.L. S.P. assisted with the expression analysis of BBS6. Transcriptome B.E.G. performed immunocytochemical analyses in cells. D.K.K., J.L. and U.W. conceptualized the studies. D.K.K. and J.L. drafted the manuscript and U.W. revised and wrote the manuscript. H.L.M. supervised S.P. and revised the manuscript.

---

**Supplementary Figure S1.**
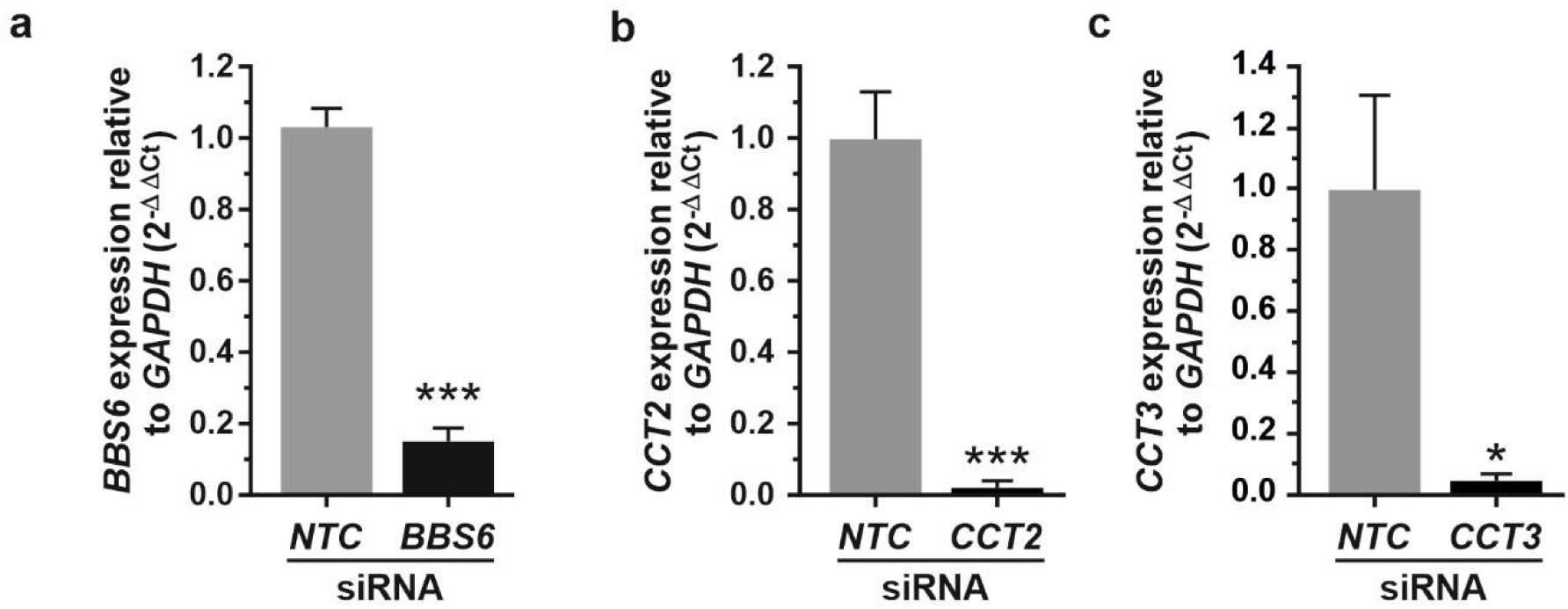
Validation of siRNA-mediated knockdown chaperonins *BBS6*, *CCT2*, and *CCT3*. (**a-c**) Analysis of mRNA expression levels after siRNA-mediated knockdown (KD) of *BBS6* (**a**), *CCT2* (**b**), and *CCT3* (**c**) compared non-targeting (NTC) siRNA controls in hTERT-RPE1 cells by quantitative real-time PCR (qRT-PCR). The expression of all three mRNAs are significantly downregulated. Statistical significance was determined by the two-tailed Student’s *t*-test (a-c): *p < 0.05, **p < 0.01, ***p < 0.005.

**Supplementary Figure S2.**
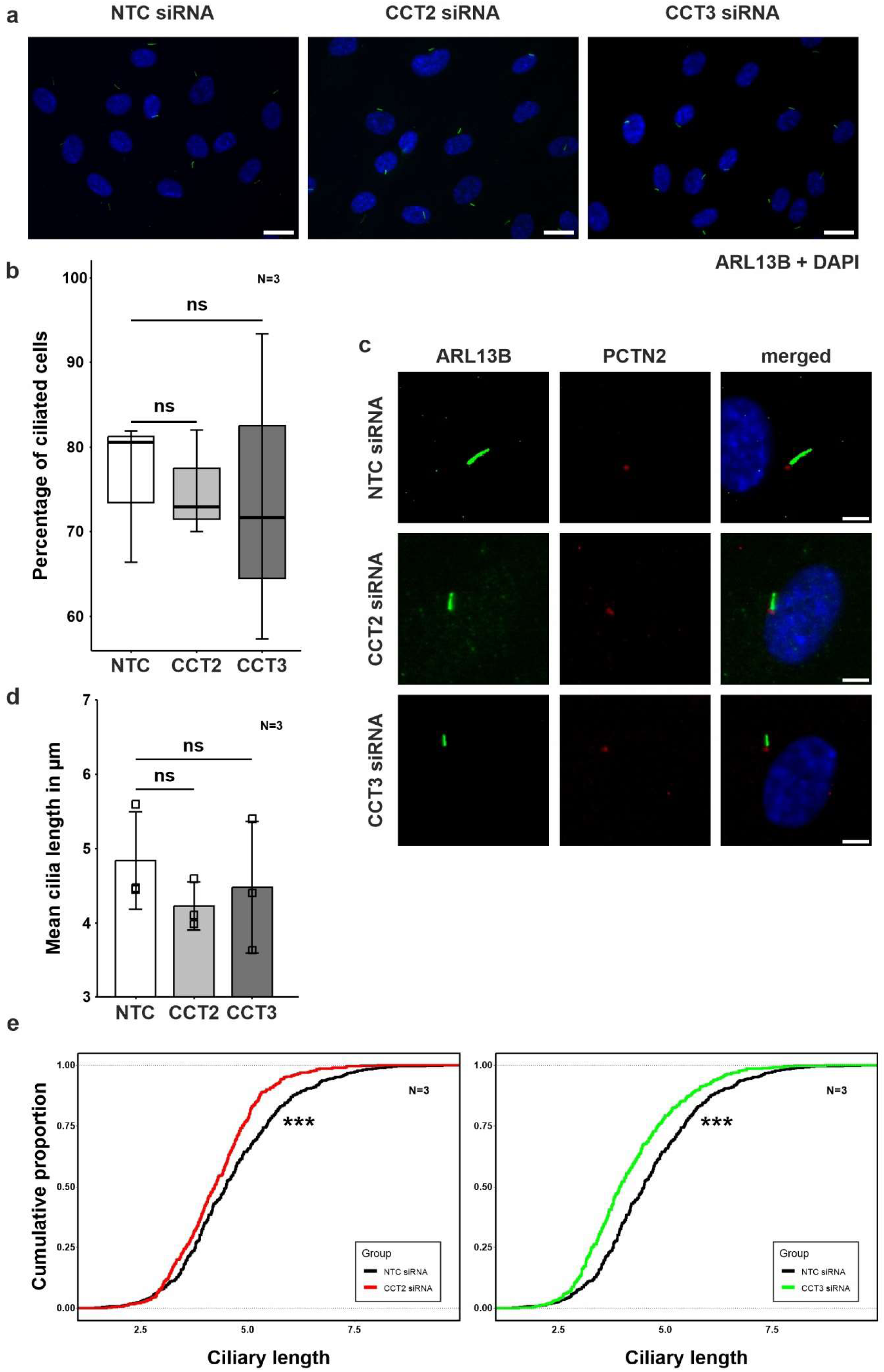
Primary cilia length is in CCT2 and CCT3 depleted hTERT-RPE1 cells. (**a**) Low magnification of immunofluorescence images of hTERT-RPE1 cells depleted for CCT2 and CCT3 by siRNA-mediated knockdown stained for the cilia marker ARL13B. (**b**) Quantification of ciliated hTERT-RPE1 cells (**c**) Immunofluorescence labelling of the cilia marker ARL13B and the ciliary base marker pericentrin 2 (PCTN2) in hTERT-RPE1 cells which were depleted for CCT2 and CCT3 by siRNA-mediated knockdown. (**d**) Quantification of mean cilia length showed no significant differences compared to NTC control. (**e**) Evaluation of the distribution of the cilia length however showed a significant reduction of long cilia in CCT2 and CCT3 depleted cells. N = number of biological replicates, n = number of analysed cells. Statistical significance was determined by the two-tailed Student’s *t*-test (b,d) and the Kolmogorov-Smirnov test (e): *p < 0.05, **p < 0.01, ***p < 0.005. Scale bar: a = 20 µm, c = 5 µm.

**Supplementary Figure S3.**
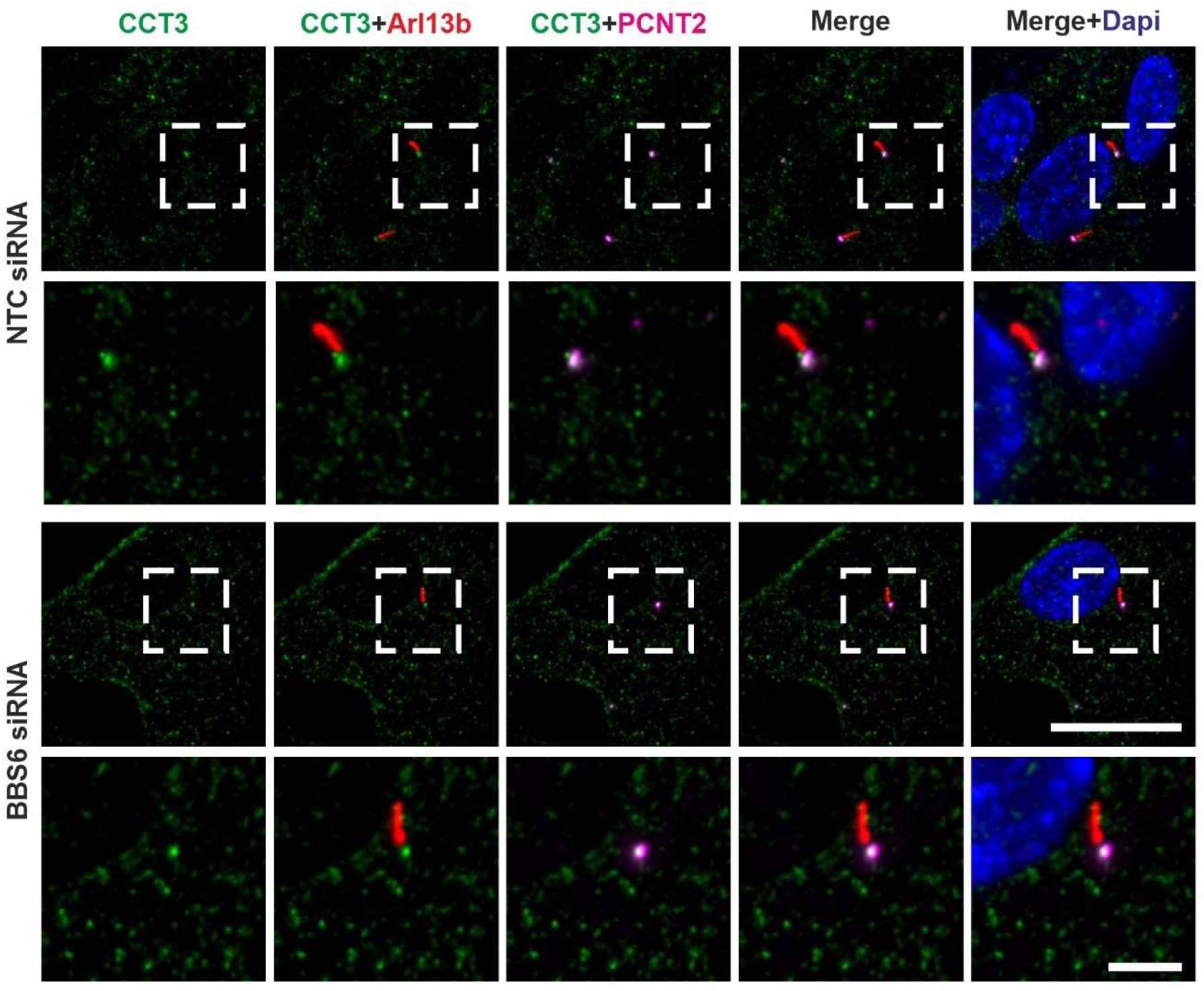
Unaltered ciliary localization of CCT3 after siRNA-mediated *BBS6* knockdown. Representative immunofluorescence images of hTERT-RPE1 cells labelled with antibodies against ARL13B as ciliary shaft marker (red), (PCNT2; magenta) as ciliary base marker, and CCT3 (green) to the base of the cilium after siRNA-mediated *BBS6* knockdown (KD) in comparison to control cells. Nuclear DNA is counterstained by DAPI (blue). The ciliary base localization of CCT3 is not altered in *BBS6* KD cells. Scale bar: 25 µm and 5 µm in magnified images.

